# Motoneurons can count: A cell intrinsic spike number memory compensates for deviations from rate coding

**DOI:** 10.1101/2025.06.16.659928

**Authors:** Lion Huthmacher, Selina Hilgert, Shana Reichert, Silvan Hürkey, Stefanie Ryglewski, Carsten Duch

## Abstract

Firing rate is an important means of encoding information in many types of neurons. A prime example is asynchronous flight as used by ∼600,000 insect species (Dudley, 2018), where wingbeat frequency and flight power output are controlled by a rate code of the flight power motoneurons (Hürkey et al., 2023). The five motoneurons that innervate the wing depressor muscle fibers translate different magnitudes of excitatory drive smoothly into changes of their common firing rates, which in turn, are linearly related to wing power output (Gordon and Dickinson, 2006). Such motoneuron input/output properties are called type-I excitability and are achieved by the expression of specific combinations of ion currents that linearize the frequency-input current curve. But are there additional motoneuron properties that compensate for acute perturbation of their rate code?

Here we combine *in vivo* electrophysiology with *Drosophila* genetics to test for mechanisms that compensate for transient perturbation of rate coding during behavior. We show that MN intrinsic properties compensate for the occurrence of extra spikes by delaying the subsequent spikes, thus restoring rate coding fidelity. The underlying mechanism is dose and phase dependent. First, compensatory increases of subsequent interspike interval durations grow with the number of supernumerous spikes that interfere with coding. Second, the magnitude of the compensation for single extra spikes depends on when during an interspike interval these occur. This mechanism depends at least in part on axonally localized HCN channels and increases the fidelity of motoneuron rate coding in the light of perturbation during flight motor behavior.

## Introduction

A century ago, Lord Adrian and colleagues (1926, 1929) demonstrated that motoneurons and many sensory neurons encode the amplitude of an input by their firing rate, which is referred to as ‘rate coding’. Although neurons may also utilize the precise timing of spikes in temporal coding schemes, as well as combinations of rate and temporal information (Ferster and Spruston, 1995; Gerstner et al., 1997), firing rate is an important means of information encoding in all brains. A rate coding neuron must respond tonically and change its firing rate proportionally to input. This is the case in neurons with a type-I frequency-input (F/I) relationship, which is characterized by low firing frequencies at small input amplitudes and smoothly increasing firing frequencies with increasing input amplitudes (Connor and Stevens, 1971a; Connor et al., 1977; Ermentrout, 1986). Type-I firing requires a tight regulation of the expression levels of specific types of voltage gated membrane channels, such as A-type potassium channels (Connor and Stevens, 1971b; Connor et al., 1977; Rush and Rinzel, 1995; Prescott et al., 2008) although it is clear that the same neuronal input/output computations can be mediated by numerous different combinations of ion channels (Marder and Goaillard JM, 2006; Marder, 2011), and computational studies indicate that the expression of multiple ion channel types with degenerate properties renders type-I firing properties more robust to variation in ion channel expression (Drion et al., 2015). In sum, numerous solutions have evolved to develop neurons that encode input amplitude with high fidelity in firing rate. However, it is less clear whether additional cell intrinsic mechanisms exist that increase the robustness of a neuronal rate code to transient perturbation.

We address this question by combining electrophysiology with neuroanatomy and genetic manipulation in *Drosophila melanogaster* flight power motoneurons (MNs) that are known to control wingbeat power output with a rate code. Flies utilize so called asynchronous indirect flight muscles (A-IFMs) for the generation of wingbeat power. Unlike other skeletal muscle fibers, A-IFMs do not contract in synchrony with MN spikes (Wilson and Wyman, 1963), but are stretch activated and form an oscillating system with their antagonists (Josephson et al., 2000) for the production of high mechanical power at fast frequencies (Böttinger and Furshpan, 1952). Although the MNs to A-IFMs fire only every ∼20^th^ to 40^th^ muscle contraction (Harcombe and Wyman, 1977), thus not controlling wingbeat on a cycle-by-cycle basis, flight MN tonic firing rate regulates wingbeat frequency indirectly by adjusting the myoplasmic calcium levels (Gordon and Dickinson, 2006). In fact, MN firing frequency is directly proportional to myoplasmic calcium as well as wingbeat frequency and amplitude, so that flight power is encoded by a MN rate code (Gordon and Dickinson, 2006; Wang et al., 2022; Hürkey et al., 2023).

The five identified A-IFM motoneurons (MN1-5, Consoulas et al., 2002) to the *Drosophila* dorsal longitudinal wing depressor muscle (DLM) respond to input with low frequency tonic firing (Ryglewski et al., 2014), exhibit linear F/I relationships, and fire tonically between ∼2 and 12 Hz during flight (Hürkey et al., 2023). MN1-5 translate common excitatory drive into common tonic firing rates, so that all 5 MNs (Harcombe and Wyman, 1977) transfer the same rate code to the different fibers of the DLM (Hürkey et al., 2023), each of which is innervated by one A-IFM MN. Similar firing rates of all 5 A-IFM MNs is behaviorally relevant and can be maintained for hours of flight (Hürkey et al., 2023). Moreover, it has previously been suggested (Harcombe and Wyman, 1977) that MN1-5 are equipped with specific properties to compensate for perturbations during ongoing rate coding. For example, during flight, sudden increases in instantaneous firing rate, as caused by evoking spikes in between subsequent spikes, lead to increases in the next interspike interval. Delaying the time to the next spike upon transient spike rate increases may restore the average target firing rate in a compensatory manner. However, neither the precise properties of this compensation, nor the underlying mechanisms are known. Here, we first confirm and quantify this phenomenon to then address the underlying mechanism. In fact, we show that transient increases in MN spiking rate are followed by compensatory firing frequency in the subsequent firing cycles. These compensatory spike rate adjustments depend on spike number and phase when the perturbation occurred. The underlying mechanism does not require CPG network connection but is MN intrinsic, and it is at least in part dependent on axonally localized HCN channels.

## Results

The firing activity of selected subsets of the five identified A-IFM motoneurons (MN1-5, Consoulas et al., 2002) that innervate the *Drosophila* dorsal longitudinal wing depressor muscle (DLM) can be recorded during tethered flight by inserting sharpened tungsten electrodes into the respective DLM target fibers (Fig. 1A, Hürkey et al., 2023). A schematic of the cell body locations of MN1-5 in the CNS and the MN axonal projections onto the 6 DLM fibers (Fig. 1B) shows that the MN1-4 somata are located ipsilaterally to their DLM target muscle and each of the 4 MNs innervates one of the DLM fibers 1-4. The MN5 soma is located contralaterally to the DLM and the MN5 axon projects to the two dorsal most DLM fibers 5 and 6 (Fig. 1B). Each muscle spike is a one-to-one reflection of the spike time of the specific MN that innervates that DLM fiber and occurs every 20^th^ to 40^th^ wingbeat (Hürkey et al., 2023). Extracellular recordings of MN spike times by the insertion of sharpened tungsten electrodes into identified DLM fibers (see methods) reveal a large amplitude spike of the muscle fiber corresponding to the spike time of the MN that innervates the inserted fiber, as well as a smaller extracellular spike that originates from the next neighboring fiber and corresponds to the spike time of the MN that innervates that fiber (see methods and Hürkey et al., 2023). Therefore, the spike times of multiple MNs can be recorded simultaneously *in vivo* during tethered flight as exemplified for MN4 and MN5 recorded from DLM fibers 5 and 4 (Fig. 1C, MN4 small spike in original trace, blue spike count; MN5, large spike in original trace, purple spike count). Confirming previous work (Hürkey et al., 2023), in all recordings of MN4/5 pairs (we recorded a total of 110 animals in this study), both MNs fire tonically and at similar frequencies, but out-of phase (Fig. 1C). During stationary tethered flight with constant and static (not-changing) sensory input, MN firing rate (Fig. 1C) and wingbeat frequency are rather constant. MNs exhibit low firing rates between ∼2 and 12 Hz encoding for wingbeat frequencies between ∼180 and 220 Hz (Hürkey et al., 2023). We now ask, what happens in response to a transient perturbation of the rate code in one MN?

**Figure 1.**
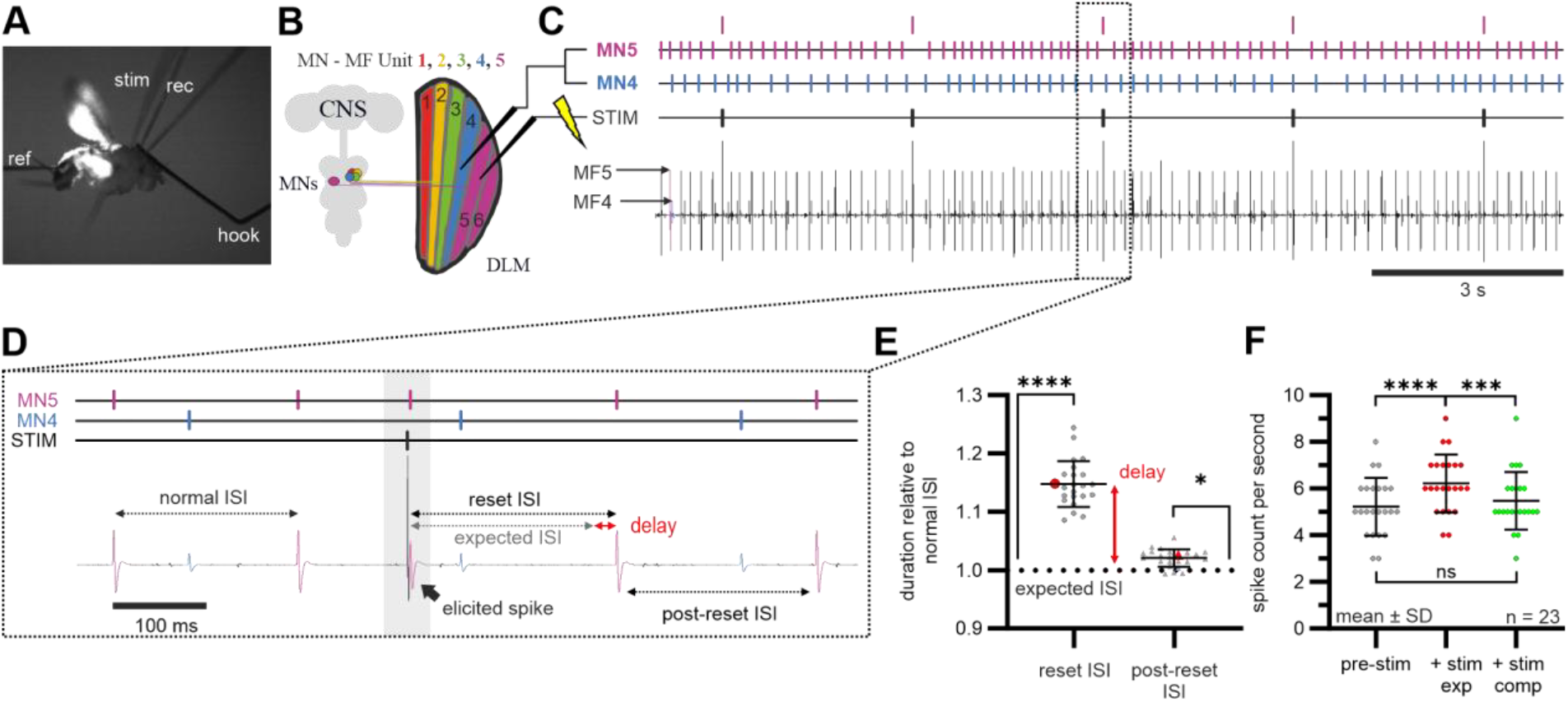
Antidromic MN stimulation resets the spike cycle and increase the subsequent ISIs. Extracellular recordings from the DLM fibers 5 and 4 during tethered flight (**A, B**) reveal (**C**) the spike times of MN5 and MN4 during flight motor behavior. Lower trace in (**C**) is the original recording of MN4 and MN5 with one electrode, upper trace the spike event trace for MN4 (blue) and MN5 (purple). In addition, antidromic stimulation (see large amplitude stimulation artifact) can be used to selectively elicit extra spikes in MN5. (**D**) The stimulation artifact is reliably followed by an antidromically elicited extra spike (see black arrow in recorded trace and purple spike time in upper trace) but not of MN4 (see small spikes in lower recorded trace and blue spike times in upper trace). An interspike interval (ISI) between subsequent MN5 spikes before the antidromic stimulation is referred to as normal ISI. The mean of the last 5 normal ISIs before the stimulation is referred to as expected ISI (gray arrow). The first ISI after the antidromic stimulus is referred to as reset ISI because it starts with the extra spike and ends with the first subsequent spike. The difference between the duration of the expected ISI and the reset ISI is referred to as spike delay (red arrow). The subsequent ISI is referred to as post-reset ISI. (**E**) Diagram shows the difference of the reset ISI and the post-reset ISI relative to the mean expected ISI (see dotted line). The first ISI (reset ISI) after an antidromically evoked extra spike was roughly 15 % larger than the mean expected ISI (Friedman test with Dunn’s post-hoc comparison, ****p < 0.0001, n = 23 animals), and the second ISI (post-reset ISI) after an antidromically evoked extra spike was roughly 2-3 % larger than the mean expected ISI (Friedman test with Dunn’s post-hoc comparison, *p = 0.0244, n = 23 animals). **(F)** Quantitative comparison of the number of MN spikes within one second before the ISI that received the extra spike (left, pre-stim) with the expected spike number during the subsequent second including the extra spike (middle, + exp) and the spike number observed within the subsequent second including the extra spike (right, + stim comp). (n = 23 animals, Friedman test with Dunn’s post-hoc comparisons, ****p < 0.0001, ***p = 0.0005 and p = 0.5535, respectively).

### Compensatory adjustments to transient perturbations of spike rate

To test this, we transiently perturbed the rate code of MN5 in wildtype animals during tethered flight by inducing an extra spike through antidromic stimulation (Fig. 1C). We repeated this perturbation every 3 seconds (Fig. 1C). A representative selective enlargement of the recording of MN5 and MN4, showing a few spikes before and after an antidromic stimulation of MN5, indicates that this perturbation affects the interspike interval (ISI) duration between subsequent MN5 spikes, but not that of MN4 (Fig. 1D). The two spikes before antidromic stimulation show the normal interspike interval (ISI) of MN5 and match an expected ISI that is calculated as the average of the last 5 ISIs before the antidromic stimulation (see methods). In the example shown, the antidromic stimulation (Fig. 1D, large stimulation artifact directly followed by an elicited extra MN5 spike, gray shaded box) occurs within the next expected ISI. The elicited extra MN5 spike is then followed by the normal expected ISI plus an extra delay. We define the first ISI after an antidromic stimulation as reset ISI, because the MN5 spiking cycle is reset to phase 0 at the time point of the extra spike so that the elicited extra spike always shifts the next orthodromic spike to a later time point than would be expected without stimulation. Moreover, the ISI between the antidromic spike and the subsequent orthodromic spike matches the expected ISI (Fig. 1D, gray arrow) plus an extra delay (Fig. 1D, red arrow). Importantly, this is carried over also to the second ISI after the induced extra spike, which we define as post-reset ISI (Fig. 1D). By contrast, the firing rate or ISI durations of MN4 are not significantly affected by antidromically induced extra spikes of MN5 (Figs. 1C, D). Calculating the median of the reset ISI durations and the post-reset ISI durations (from >75 antidromic stimulations per animal) for 23 animals and relating these values to the expected ISI duration without stimulation (Fig. 1E, dotted line), reveals that just one extra spike causes a roughly 15% and highly significant increase in the reset ISI duration (Fig. 1E, ****, p < 0.0001, Friedman test with Dunn’s post-hoc comparison) directly after the antidromic spike. In addition, the extra spike also has a small (∼2%) but statistically significant (Fig. 1E, *, p = 0.0244, Friedman test with Dunn’s post-hoc comparison) effect on the next ISI (post-reset ISI). The median values of the example recording shown in figures 1C, D are colored red and are in line with the variance of the medians derived from 23 animals (Fig. 1E).

Do the delays that are caused after extra spikes compensate for the transient increases in firing rate as caused by evoked extra spikes? To test this, we count the numbers of MN spikes that occur within the second before the stimulated ISI (Fig. 1F, left) and compare this with (i) an expected spike number including the stimulation (equaling the spike number in the second without stimulation plus the evoked extra spike, Fig. 1F, middle) as well as with (ii) the observed spike number within the second including the stimulation (Fig. 1F, right, for details see methods).

The data show that without any compensatory adjustment, the extra spike would cause a significant increase in spike number per second (Fig. 1F, compare left and middle, Friedman test with Dunn’s post-hoc comparison, ****p < 0.0001). However, the observed spike numbers including the extra spike are significantly lower than the expectation (***p < 0.0005), but are not significantly different from the MN spike counts in the same period of time before the stimulation (Fig. 1F, p = 0.5535). Therefore, single extra spikes can be fully compensated for by the prolongation of the reset and the post-reset ISI, thus increasing the fidelity of the rate code.

### Phase dependency of compensatory spike rate adjustments

We next tested whether the prolongation of the reset ISI (spike delay, Figs. 1D, E) depends on when during the expected ISI an extra spike is elicited. First, for one representative animal, the relative amplitudes of the reset ISIs are plotted as a function of the phase when the stimulation occurred within two subsequent spikes (Fig. 2A, gray dots), with phase 0 depicting the time point of the last spike before the antidromic stimulus and phase 1 the time point of the next expected spike. The reset ISI duration is plotted relative to the mean expected ISI duration (derived from the last 5 ISIs before the stimulation, dotted line). The data indicate that increases in the reset ISI are larger the closer the extra spike is elicited to the last regular spike (Fig. 2A). To compare the effect of phase across animals, phase within the expected ISI is divided into 4 bins (first, second, third, and fourth quarter of the expected ISI) and for these 4 bins, median values for the resulting reset ISI durations are calculated from the 332 single stimuli given every 3 seconds in this animal (Fig. 2A, blue dots). To rigorously quantify the effect of phase this is then done for 23 animals (Fig. 2Ai). The mean values for the four bins of the representative animal shown in figure 2A are plotted in blue to show that these are right within the data variance that is observed across all 23 wildtype animals tested (Fig. 2Ai, gray dots). The data clearly reveal a highly significant phase dependency, because the increase in the duration of the reset-ISI relative to the mean expected ISI is larger the closer the extra spike occurs after that last regular MN5 spike. The reset ISI is increased by ∼30% in the first bin, by ∼20% in the second bin, by ∼10% in the third bin, and by roughly 5% in the fourth bin (Fig. 2A). Each phase bin differs significantly from the next one (Fig. 2Ai, RM one-way ANOVA with Sidak’s post-hoc comparisons,****p < 0.0001, ****p < 0.0001, **p = 0.0051, respectively), indicating that the temporal resolution of the phase dependency is higher than 25% of the phase, which equals roughly 50 ms at the average MN firing rate of 5.2 ± 1.3 Hz (mean and SD) that we find in CS controls. In sum, the closer an extra spike occurs to previous endogenous spike, the higher the increase in instantaneous firing rate at this moment, the larger the increase in the reset ISI, and thus the larger the compensation for the transient increase in firing rate.

**Figure 2.**
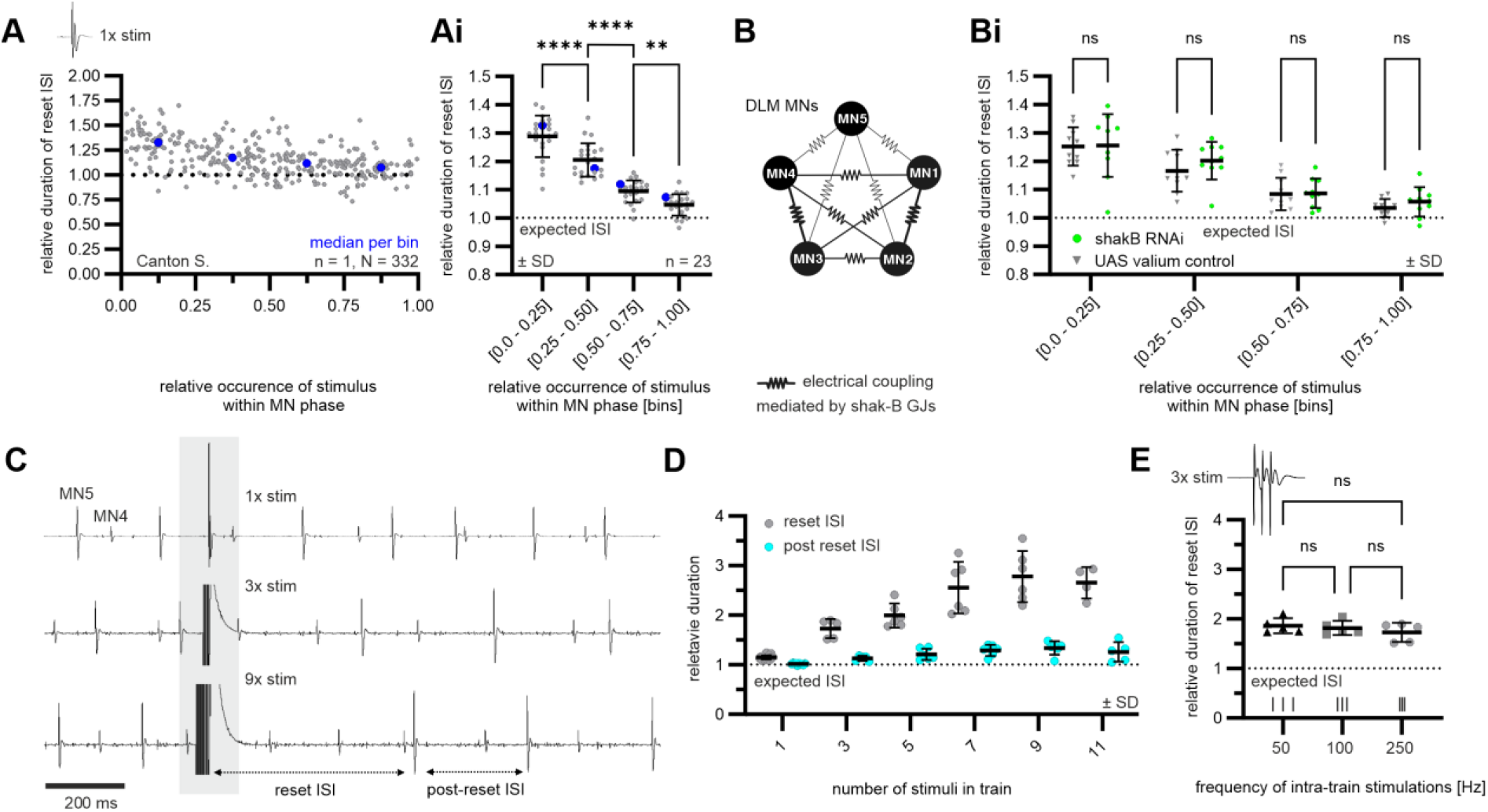
MN spike reset ISI duration is cell-intrinsic, timing- and dosage-dependent and frequency-independent. (**A, Ai**) The relative duration of the reset ISI depends on the timing of the resetting stimulus within each perturbed MN phase. (**A**) Representative scatter plot of all reset ISI durations of one wildtype MN recording resulting from repeated single pulse stimulations administered at different phases of MN5 spiking. Median reset ISI duration calculated for each quartile bin of the MN phase are shown as blue dots and correspond to those in (**Ai**). (**Ai**) Reset ISI durations for stimulations occurring in the first to last quartile of the MN phase are highly significantly reduced between bins (Fig. 2Ai, RM one-way ANOVA with Sidak’s post-hoc comparisons,****p < 0.0001, ****p < 0.0001, **p = 0.0051, respectively). (**B, Bi**) Knockdown of gap junction protein shakB does not affect the relative duration of the reset ISI compared to genetic control (**Bi**), indicating no involvement of the electrically coupled network of the DLM MNs (**B**) in generating the reset ISI elongation (Fig. 2Bi, Ordinary two-way ANOVA with Tukey’s post-hoc comparisons, left to right: p = 0.9070, p = 0.2396, p = 0.9268, p = 0.4636). (**C, D**) The duration of the MN5 reset ISI increases relative to the count of stimulations administered in train. (**C**) Representative recordings for one, three and nine stimulations in train. (**D**) Post-reset ISI duration also increases for increased counts of stimulations in train. (**E**) Reset ISI duration does not depend on the intra-train pulse frequency in triple pulse stimulations (Kruskal-Wallis test with Dunn’s post-hoc comparisons, each p > 0.9999).

### Compensatory spike rate adjustments are caused by MN intrinsic mechanisms

We have previously found that the 5 MNs are reciprocally connected by weak electrical synapses (Fig. 2B) which organize the phase relationships between MN1-5 during flight motor pattern generation (Hürkey et al., 2023).

Therefore, it is possible that the delay of the spikes following an antidromically evoked extra spike is an indirect consequence of the weak electrical coupling in the motoneuronal CPG network. However, targeting UAS-ShakB RNAi exclusively to MN1-5 does neither affect the durations nor the phase dependencies of the reset ISI as compared to the corresponding genetic control (Fig. 2Bi, Ordinary two-way ANOVA with Tukey’s post-hoc comparisons, left to right: p = 0.9070, p = 0.2396, p = 0.9268, p = 0.4636), or as compared to wildtype flies (compare Figs. 2Ai and 2B). The effectiveness of the RNAi knock-down for ShakingB encoded electrical synapses in MN1-5 has previously been confirmed (Hürkey et al., 2023). These data are consistent with MN intrinsic mechanisms to underlay compensatory spike rate adjustments.

### Compensatory spike rate adjustments occur in a dose dependent manner

We started with single antidromically evoked extra spikes during normal in-flight tonic MN firing. However, real live disturbances, such as a brief gust of side wind, may cause the transient occurrence of multiple extra spikes. Therefore, we next tested whether the increases in reset ISI, which constitutes the amplitude of the compensatory spike rate adjustment, is not only phase dependent (Fig. 2Ai), but also dose dependent. In fact, short bursts of 3 extra spikes evoked in MN5 at 250 Hz during tethered flight increased the reset ISI substantially more than a single evoked extra spike (Fig. 2C, top two traces). Increasing the number of extra spikes from 3 to 9 in the same recording prolonged the MN5 reset ISI even further, so that 3 cycles of MN4 firing now fit into one MN5 reset ISI (Fig. 2C, lower trace). Quantification reveals that the reset ISI increased from 1.3-fold the normal expected ISI to nearly 3-fold the normal ISI, but is not further increased with 11 extra spikes (Fig. 2D, gray dots). Therefore, for extra spikes that occur at 250 Hz during a normal ISI, the next ISI (reset ISI) is prolonged in a dose-dependent manner (Fig. 2D, gray dots). The underlying mechanism covers a working range from 1 to 9 spikes presented in a short burst but saturates at larger spike numbers per burst. Given that we also observed a slight increase in the duration of the post-reset ISI (Fig. 1E, the second MN5 spiking cycle after the antidromically evoked extra spike) this is quantified in the same animals. The post-reset ISI increases with the number of extra spikes and saturates at roughly 10 extra spikes, but the maximum amplitude of the duration increase is much smaller (∼40%) for the post-reset ISI (Fig. 2D, cyan dots) as compared to ∼300% for the reset ISI (Fig. 2D, gray dots). To test whether the number or the frequency of the transiently evoked extra spikes are reflected in the increases in reset ISI duration, we stimulated with 3 extra spikes within a normal ISI during in-flight MN5 tonic firing, but presented these either at 50 Hz, at 100 Hz, or at 250 Hz (Fig. 2E). The data clearly demonstrate that it is the number (Fig. 2D) but not the frequency of extra spikes (Fig. 2E, Kruskal-Wallis test with Dunn’s post-hoc comparisons, each p > 0.9999) that is encoded in the increased duration of the reset ISI. Therefore, we refer to the MN intrinsic compensatory adjustments in firing rate as a spike number memory.

### HCN channel as a candidate player in the mechanism for a spike number memory

Possible mechanisms underlying a MN intrinsic spike number memory include some kind of activity dependent modulation of existing membrane currents that affect firing rate, or membrane channels that affect spike rate and have slow kinetics that operate at the required time courses of ∼100 to 600 ms that are relevant to affect MN firing between 2 and 12 Hz. We know that adult flight MNs express the A-type potassium channels Shaker (K_v_1 homolog) and Shal (Kv4 homolog) (Ryglewski and Duch, 2009; Werner et al., 2020), the delayed rectifier potassium channels Shab and Shaw (Ryglewski and Duch, 2009; Kilo et al., 2014; Werner et al., 2020) as well as the Ca_v_2 and Ca_v_3 VGCC homologs cacophony (Dmca1A) and DmαG (Ryglewski et al., 2012; Bell et al., 2025) and BK calcium activated potassium channels. However, none of these channels have sufficiently slow kinetics to be a likely candidate for affecting MN spiking at time courses between 100 and 600 ms, although we cannot exclude the participation of these channels in the spike number memory. However, hyperpolarization activated, cyclic nucleotide gated channels seem strong candidates because they can have slow activation and inactivation kinetics and are modulated by cyclic nucleotides that can in turn be regulated by changes in firing rate (Biel et al. 2008). HCN channels have been reported functionally important in gustatory (Lee et al., 2025) and mechanosensory receptor neurons (Trombley et al., 2023) as well as at the presynaptic terminal of larval MNs (Hegle et al., 2017). Although HCN mutants show adult motor phenotypes, HCN channel expression in adult flight MNs has previously not been demonstrated. We show that ZD7288, a potent blocker of cardiac and neuronal HCN channels, causes a significant hyperpolarization of the MN5 membrane potential as recorded from the soma in control animals (Fig. 3A, Kruskal-Wallis test with Dunn’s post-hoc comparisons, **p = 0.0014). By contrast, HCN mutants show no voltage response to ZD7288 bath application whereas controls respond with a hyperpolarization of 10 mV on average, which is a highly significant difference (Fig. 3B, two-tailed Mann-Whitney test, ***p = 0.0002). This is consistent with a fraction of HCN channels open near resting membrane potentials. Blockade with ZD7288 reduces membrane conductance for monovalent cations. Near rest, changes in potassium conductance have little effect, but the decreased sodium conductance causes a hyperpolarization (Biel et al. 2008). Acute blockade of HCN also causes a significant reduction of the tonic firing frequency of adult MNs in response to somatic square pulse current injections (Figs. 3C, D), indicating that HCN channels affect MN firing rate. However, in HCN mutants, MN firing rate and wingbeat frequency as observed during tethered flight is not significantly different from wildtype control or the controls that were used for ShakingB knock down flies (Fig. 3E, Brown-Forsythe and Welch ANOVA with Dunnett’s T3 post-hoc comparisons, left to right: p = 0.9843, p = 0.1483, p = 0.1548). Moreover, somatic current clamp recordings do neither reveal any sag potentials as typical for neurons with somatically localized HCN channels (Biel et al. 2008), nor can h-current be recorded in response to hyperpolarizing command potentials applied to the soma in voltage clamp.

**Figure 3.**
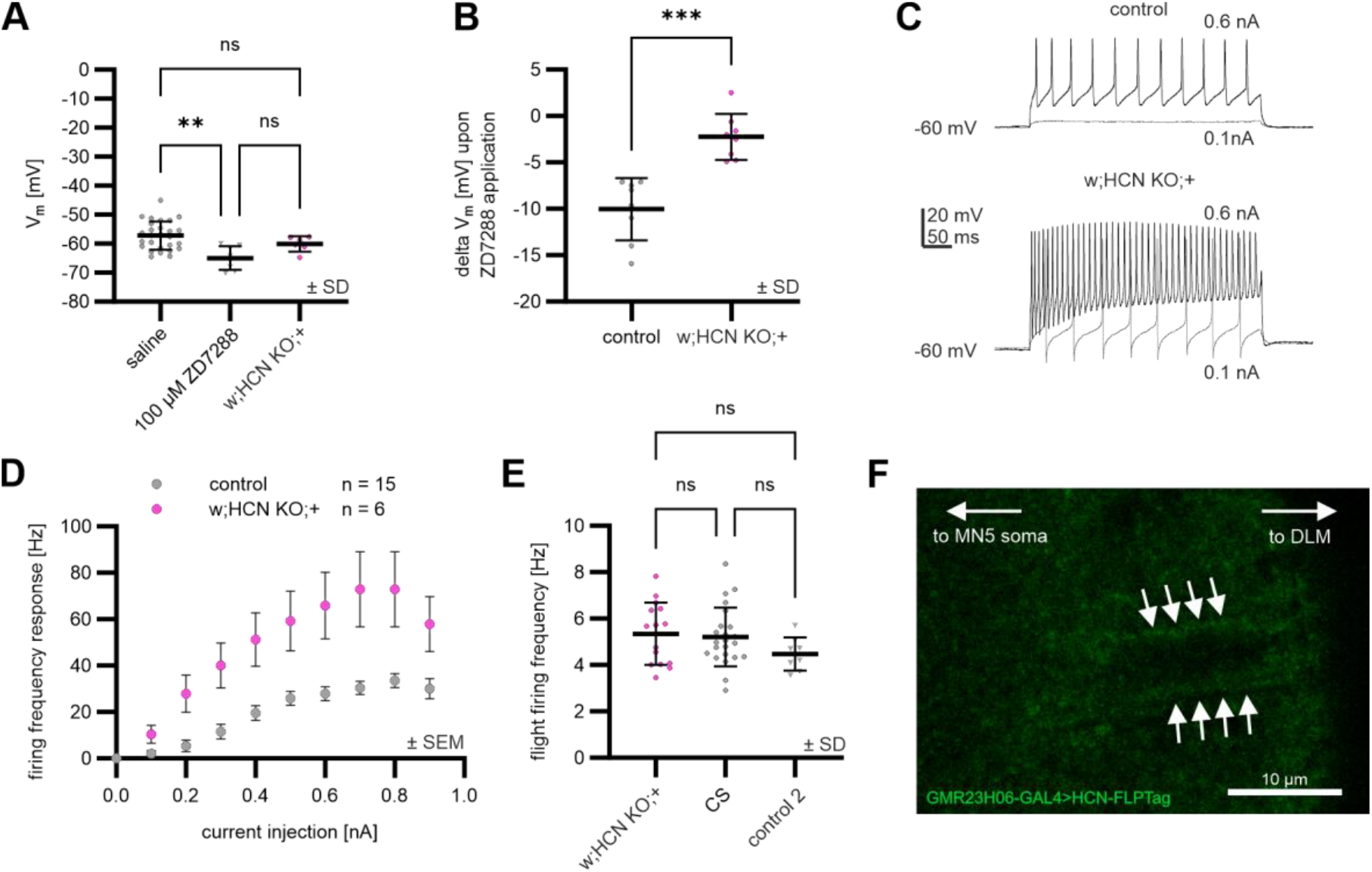
Effects of pharmacological and genetic abolishment of Ih on MN5 membrane properties. (**A**) Resting membrane potential (Vm) in control animals under control conditions, after application of ZD7288, and in HCN channel mutants under control conditions (Kruskal-Wallis test with Dunn’s post-hoc comparisons, left to right: **p = 0.0014, p = 0.6675, p = 0.3455). (**B**) Change of Vm upon application of specific Ih blocker ZD7288 in control animals and in HCN mutants (Two-tailed Mann-Whitney test, ***p = 0.0002). (**C**) Example traces of MN5 membrane excitability after 400 ms lasting current injection of 0.6 nA and 1 nA for control animals and HCN mutants. (**D**) Excitability of MN5 under control conditions expressed as firing frequency upon 400 ms lasting current injection between 0 and 0.9 nA in control and Ih mutant animals. (**E**) No significant difference between in-flight firing frequency of HCN KO animals compared to CS wildtype and the controls that were used for ShakingB knock down flies (Brown-Forsythe and Welch ANOVA with Dunnett’s T3 post-hoc comparisons, left, right, up: p = 0.9843, p = 0.1483, p = 0.1548) (**F**) Localization of HCN channels labeled by GFP Flip Tag along DLM MN axons (white arrows) within the posterior dorsal mesothoracic nerve.

These observations are consistent with HCN channels being localized in distal axonal or dendritic compartments that are hidden to somatic voltage clamp control in these large neurons with their spike initiating zone at a distance of several hundred micrometers from the soma (Kuehn and Duch, 2013), >6000 µm total dendritic length (Duch et al., 2008), and >4000 dendritic branches (Vonhoff and Duch, 2010; Ryglewski et al., 2017). In agreement with this, immunohistochemical data on *Drosophila* HCN channels that are endogenously tagged with GFP suggest localization along the axons of flight MNs close to the spike initiating zone at the nerve root (Fig. 3F). These data suggest that adult flight MNs express functional HCN channels in neuronal compartments that are distal to the soma, likely including the axon, but potentially also dendrites. Moreover, a fraction of the MN HCN channels is likely open near resting membrane potential and HCN channel expression affects tonic MN firing responses.

### HCN channels contribute to the spike number memory of adult flight MNs

In HCN null mutants (see methods) the increase in ISI duration after an extra spike is significantly reduced, but not eliminated. The phase dependency of the spike number compensation remains, but the effect is significantly smaller in all 4 phase bins (Fig. 4A, Ordinary two-way ANOVA with Tukey’s post-hoc comparisons, left to right: ****p < 0.0001, ***p = 0.0009, *p = 0.0403, *p = 0.0221). Similarly, the larger spike number compensation that is seen upon delivering 3 extra spikes in one ISI (Figs. 2C, D) is highly significantly reduced in HCN channel mutants (Fig. 4B, Ordinary two-way ANOVA with Sidak’s post-hoc comparisons, ****p < 0.0001). However, the small but significant increase of the post-reset interval that is induced by 3 stimuli within one ISI (Figs. 2C, D) is not significantly affected in the absence of HCN channels (Fig. 4B, p = 0.9754). These data are consistent with a function of HCN channels to significantly increase the duration of the reset ISI. Given that the effect of the extra spike is not eliminated, HCN channels must act in concert with additional mechanisms.

**Figure 4.**
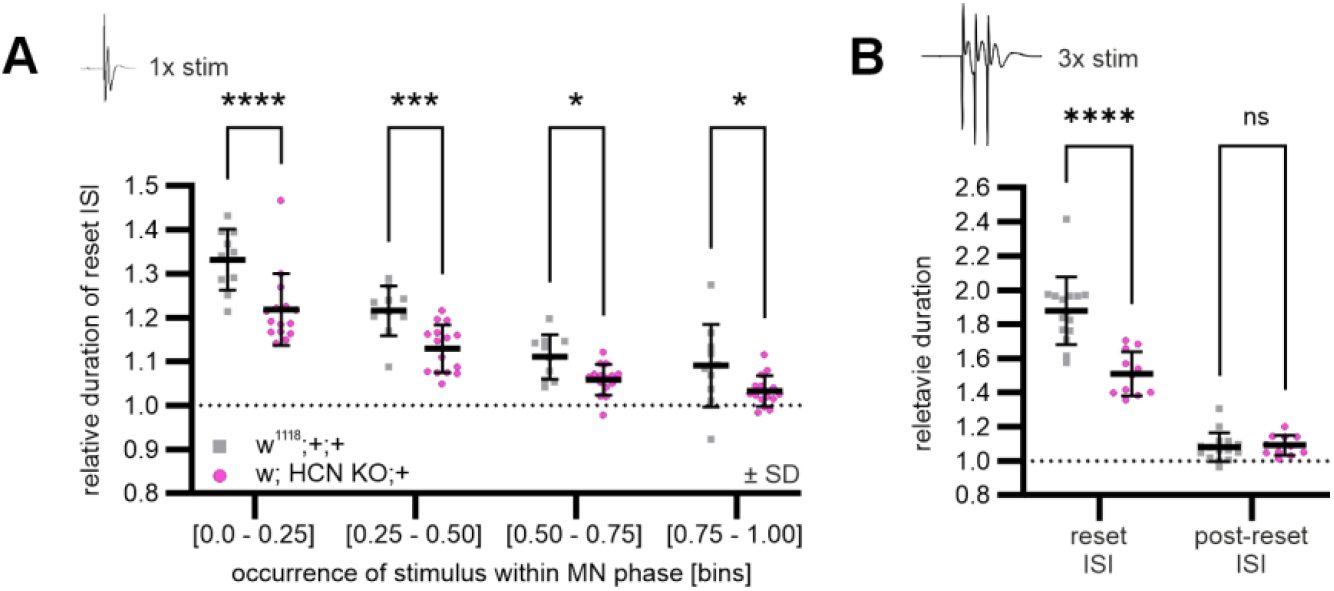
HCN channels contribute significantly to the compensation for extra spikes. (**A**) The relative duration of the MN5 reset ISIs resulting from single pulse stimulations are significantly decreased in HCN mutant flies compared to control (w^1118^). Please note that the w1118 control shows no significant differences in the reset ISI durations as compared to Canton S controls (compare Fig. 2Ai). This effect is independent of the bin of the occurrence of the stimulus within the MN phase (Ordinary two-way ANOVA with Tukey’s post-hoc comparisons, left to right: ****p < 0.0001, ***p = 0.0009, *p = 0.0403, *p = 0.0221). (**B**) Relative duration of the MN5 reset ISI resulting from triple pulse stimulation trains is highly significantly decreased in HCN mutant flies compared to genetic controls while post-reset ISI duration remains unchanged (Ordinary two-way ANOVA with Sidak’s post-hoc comparisons, ****p < 0.0001, p = 0.9754, respectively).

## Discussion

In this study, we tested whether and how transient perturbations of a neuron’s rate code influence their subsequent spiking rate. We find that the occurrence of supernumerous spikes is compensated for by a cell-intrinsic, phase- and dose-dependent mechanism that reduces the instantaneous firing rate to restore overall firing rate and thus support coding fidelity. Instantaneous firing rate is transiently lowered by resetting the spike cycle to the time point of the evoked extra spike and by prolonging the durations of the next ISI (reset-ISI) and, to a lesser degree, also the ISI after that (post-rest ISI). The principal phenomenon has been observed in multiple species, but we characterize it in greater detail, quantify the degree of compensation to assess its physiological relevance, and we begin to reveal the underlying mechanism.

### Spike rate compensation by resetting and prolonging the spike cycle

A tonically firing neuron can either passively follow a rate that originates from other sources, or it can produce the rate code by endogenously translating the sum of many postsynaptic potentials into a tonic firing frequency. The latter is the case for the *Drosophila* flight MNs studied here (Hürkey et al., 2023) that exhibit classical type-I excitability. Similar to early experiments in crayfish (Preston and Kennedy, 1962), locust (Wilson, 1964), and *Drosophila* (Levine, 1973; Harcombe and Wyman, 1977; Koenig and Ikeda, 1983) our data show that in tonically discharging MNs, evoked extra spikes reset the spike cycle. Resetting is achieved by delaying the next spike by approximately one normal interspike interval. Importantly, resetting occurs in response to both orthodromic spikes as evoked by somatic current injection (Koenig and Ikeda, 1983) and antidromic spikes (Harcombe and Wyman, 1977). This indicates that resetting of the spiking cycle is a direct consequence of spike invasion of the spike initiating zone. Our data further extend previous findings that the firing cycle reset is followed by an average interspike interval (ISI) plus an additional delay (Harcombe and Wyman, 1977; Koenig and Ikeda, 1983). The duration of the extra delay is tuned in a phase dependent manner. In addition, we now provide quantitative evidence that this extra delay provides a significant and reliable compensation of the increase in firing rate caused by extra spikes. Furthermore, this compensation is not limited to single extra spikes but increases in magnitude with the number of supernumerous spikes. In fact, a range of 1 to 9 supernumerous spikes is counteracted by dose-dependent increases in ISI. The dose is defined by the number of extra spikes rather than by their frequency. Finally, prolonging is not restricted to the first ISI after the extra spike (reset ISI) but carries over into the subsequent ISI, although at a strongly reduced magnitude. In sum, our data are consistent with a MN intrinsic spike number counter that is employed to compensate for sudden increases in firing rate and thus aids to maintaining rate coding fidelity upon perturbation.

### Physiological relevance of MN spike rate compensation

Maintaining constant tonic MN firing rates is essential for the control of flight power and thus behaviorally relevant (Hürkey et al., 2023). Within and across animals, all 5 DLM MNs share the same instantaneous firing rates and tune these in common over time to adjust wingbeat frequencies to changing power demands. Mechanistically, common firing rates of all MNs are achieved by combining qualitatively and quantitatively similar MN input/output properties (all 5 MNs show nearly identical type-I firing response-frequency / current input (F/I) relationships) with common synaptic drive to the MNs (Hürkey et al., 2023). In fact, connectomics suggest that the DLM MNs share nearly 90% of their excitatory input from that same 129 presynaptic cholinergic partners (Takemura et al., 2023). It is thus plausible that all DLM MNs typically experience sudden perturbations of their rate code in unison. For example, imitating wind gusts, as may be occurring during normal flight, by delivering brief air pulses during tethered flight can in fact cause transient periods of supernumerous spikes across all MNs. Spike resetting and prolonging the subsequent ISI may stabilize power output by restoring the target firing rate over time. Please note that in asynchronous flight, the MNs do not control muscle contractions on a wingbeat-to-wingbeat basis (Gordon and Dickinson, 2006), but the MNs fire only every 20^th^ to 40^th^ wingbeat, so that the translation of changes in MN firing rate into wingbeat frequency is slow. This acts like a low pass filter that makes the effects of transient increases in MN firing rate less severe but likely aids the efficiency of spike rate compensatory mechanisms as described here.

### Proposed underlying mechanism

Although we have not yet fully unraveled the mechanisms underlying compensatory responses to transient increases in MN firing rate, our data allow inferring key features of the phenomenon. First, the underlying mechanism is cell intrinsic because network connections within the CPG are not required. Second, the resetting mechanism must cover time frames between a few hundreds of milliseconds and a few seconds, because multiple extra spikes can delay the next spike by seconds. Third, the mechanism is initiated by extra spikes in a dose dependent manner. Forth, delaying the initiation of subsequent spikes while continuously receiving the same tonic excitatory drive that translates into regular tonic firing at a given rate can only be achieved by reducing the sensitivity to a given synaptic drive, or by increasing the duration to spike initiation that follows a given synaptic input. In theory, both could be implemented by multiple different combinations of ionic conductances. However, the properties of HCN channel match the features, and HCN channels are expressed in DLM MNs. First, h-current kinetics are slow enough to cover the time frame required. Second, h-current amplitude and kinetics can be modulated by cyclic nucleotides, which in turn, can serve as second messengers to read out spike number and provide the dose-dependency of the mechanism.

Third, our data demonstrate that a fraction of the HCN channels are open at rest and affect MN firing responses to a given excitatory input. Acute blockade of h-current increases MN firing responses to current injection, which is in line with more h-current delaying the next spike. The idea is that a transient increase in firing rate will increase cyclic nucleotide levels and thus also increase h-current and thus an increased ISI. Forth, our electrophysiology indicates that HCN channels are expressed in distal neuronal compartments, and axonal localization is also in agreement with our assessment of GFP tagged channel localization. Together these data make axonal HCN channels strong candidates for increasing the duration to spike initiation during a given synaptic drive. However, we cannot exclude additional dendritic HCN localization. In that case, dendritic input resistance would be decreased by increasing h-current conductances through the effects of cyclic nucleotides, which in turn would reduce the MN’s sensitivity to a given synaptic drive and thus also delay the next spike. Although, at current it remains unclear whether and how much dendritic or axonal HCN channels, or both, underly compensatory adjustments of MN spike rate, our data clearly show a prominent role of HNC channels, because a significant fraction of the delay to the next spike is eliminated in HCN mutants.

In sum, given that spike resetting has been described in tonically firing neurons of different species, both, the cell intrinsic mechanisms for spike counting as well as for compensatory delays of subsequent spikes as described in this study might be a common feature and provide general means for increasing the fidelity of rate coding in the CNS.

## Acknowledgments

We thank Joshua Klein (JGU Mainz) for his help with extracellular recordings, Dr. Olaf Vef (JGU Mainz) for his invaluable expert help with fly genetics and husbandry. We gratefully acknowledge the support by the German Research Foundation to CD (Du 331/15-1) and SR (RY 117/4-1), both as part of the Research Unit *RobustCircuit* (FOR 5289).

## Methods

### Animals

*Drosophila melanogaster* stocks and crosses were reared at 25°C and 60% humidity in 12h-12h light-dark cycles in transparent plastic vials (Kisker Biotech, 95 mm x 25 mm) each containing standard cornmeal food consisting of (per 100 vials): 920 ml dH_2_O, 110 g glucose, 50 g cornmeal, 10 g agar, 30 g dry yeast, 10 ml tegosept-solution (100 g/l in EtOH) and 0.5 g ascorbic acid.

Wildtype control experiments were performed on Canton Special (CS) flies. To target UAS-transgenes selectively to DLM MN1-5, we used a split GAL4 driver line (BDSC, #602182, Hürkey *et a*l. 2023) that expresses the GAL4 activating domain (AD) under control of the *anachronism* enhancer fragment GMR23H06 and the GAL4 DNA binding domain (DBD) under control of the *Netrin-A* enhancer fragment GMR30A07: w[*];P{y[+t7.7]w[+mC]=R23H06-p65.AD}attP40; P{y[+t7.7] w[+mC]=R30A07-GAL4.DBD}attP2. To specifically knock down all *shaking B* isoforms in MN1-5 this split GAL4 driver line was crossed to UAS-shakB RNAi flies (BDSC, #57706): y[1]sc[*]v[1]sev[21];+;P{y[+t7.7]v[+t1.8]=TRiP.HMC04895}attP2 and to empty UAS-VALIUM (BDSC, #35786) as control: y[1]v[1];+;P{UAS-GFP.VALIUM10}attP2.

### Generation of the HCN (Ih) excision mutant

Excision of ∼3 kb of the Ih gene was achieved by flipase mediated excision of DNA marked by FRT sites. Excision was initiated between PBac{w[+mC]=WH}Ih[f03355] (Bloomington #85660, RRID:BDSC_85660) and PBac{w[+mC]=RB}Ih[e01599] (Bloomington #17970, RRID:BDSC_17970; Thibault et al., 2004) in the presence of hsFLP^1^. Excision was confirmed with PCR with an expected band size of 1.7 kb if the deletion occurred. Forward primer 5’-TGCATTTGCCTTTCGCCTTAT-3’ and reverse primer 5’-AATGATTCGCAGTGGAAGGCT-3’ were used (Parks et al., 2004).

For control experiments w[1118];+;+ were used.

### Electromyography/extracellular recording of MN spike-times

Spike times of DLM MNs during tethered flight behavior were recorded extracellularly via the electrical activity of their target muscle fiber(s). MN1-4 each exclusively innervate one DLM fiber (MF1-4) and MN5 jointly innervates MF5 and MF6. Each DLM MN action potential results in one large, transient postsynaptic depolarization of its target DLM fiber which allows to directly infere the respective MN spike time.

In preparation of the EMG recordings, one to three days old male flies were cold-anesthesized on ice for 10 s and then transferred to a metal cooling plate (3 - 6 °C). A metal hook (0.1 mm tungsten wire clamped to a 5 cm carbon rod) was glued between head and thorax using UV Glass Adhesive (Super Glue Corporation) which was cured under ultraviolet light (400 - 500 nm, Mega Physik Dental Cromalux-E Halogen Curing Light Unit) for 60 s. Flies were then allowed to recover from anaesthesia for 10 mins.

The tethered flies were then suspended and positioned inside the recording-setup via a micro-manipulator (Sutter, MP225). Additional micro-manipulators were used to position the reference-, recording- and stimulation-electrode (sharpened 0.1 mm tungsten wire) necessary for EMG recordings as well as a red laser light barrier to record the wingbeat. The reference electrode was inserted dorsally between the fourth and fifth abdominal segment. The recording electrode was inserted dorsally between the left anterior dorsocentral bristle and the mid-sagittal plane through MF5/6 into MF4 to extracellularly record MN5 and MN4 spike times. Insertion at this position guarantees the first MF spike signals that appear to stem from MF5/6 and the second spike signals upon deeper insertion to stem from MF4. Signal amplitude increases with insertion depth of the electrode into the MF which allows for unambiguous identification of large amplitude spikes to stem from MF5/6 and small spikes to stem from MF4. The stimulation electrode was exclusively inserted into MF5/6 at the third acrostichial hair following the anterior dorso-central bristle in anterior direction.

### Antidromic MN stimulation

Manipulation of MN5 spike times during tethered flight was achieved by electrical single-pulse or pulse-train stimulation of MF 5/6. In experiments of the first case (Fig. 1; Fig. 2 A, Ai, Bi; Fig. 4B), single square pulse stimulations of 0.25 ms duration were administered every three seconds to MF5/6. The voltage was adjusted from an initial 5 V towards 10 V until each stimulation pulse reliably triggered an additional MF spike occurring approximately 1 ms after stimulation. Triggering of such an additional MF spike within the current phase of MN5 spiking delays the next expected spike by approximately one full cycle of the current average cycle duration when measured from the triggered MF spike. Stimulation of single MF spikes therefore *resets* the current cycle of MN5. This indicates membrane potential alterations at the MN5 spike initiating zone resulting from initial axon terminal stimulation and stimulus propagation along the antidromic direction. In experiments of the second case, following successful establishment of single-pulse MN resetting, stimulations were administered as trains of 0.25 ms square pulses either as three pulse trains of 50 Hz, 100 Hz or 250 Hz inter-pulse frequency (Fig. 2 E) every 5 seconds or as 3 to 11 pulse trains of 250 Hz inter-pulse frequency (Fig. 2 C & D; Fig. 3 H) every 10 seconds.

### Data acquisition

Analog MF recordings were amplified 100x by a Differential AC Amplifier (Model 1700, A-M Systems), low-pass filtered at 100 Hz and high-pass filtered at 500 Hz and digitized by the Digidata 1550B (Molecular Devices) at a sampling rate of 20 kHz. The signal of the laser light barrier was not amplified and digitized directly. 5 to 10 V square pulse stimulations of 0.25 ms duration were generated by a Grass Medical Instruments S48 G stimulator, isolated by a Grass Medical Instruments SIU5B stimulus isolation unit and administered to the stimulation electrode (−) and ground (+). Recordings were captured using the software AxoScope V. 10.7.0.3 (Molecular Devices, 2016) and saved as .abf-files.

Starting the data acquisition, MF5/6 and MF4 spikes and the wingbeat signal were recorded for 120 s as unmanipulated control flight. Subsequently one of the stimulation protocols was applied.

### Data analysis of MN spike times

For data analysis, recorded .abf-files were imported to Spike2 (v.7.2.). Semi-automatic spike sorting was performed via template-matching of spike shapes using the Spike2 *Wavemark* function and subsequent manual proof reading. Occasional interference of MF5/6, MF4 or stimulation artefact spikes within the same recording trace were resolved using the *Split Spike* function within the *Edit Wavemark* funtion. Spike times of MF5/6, MF4 and the stimulus pulses were subsequently exported as .txt files and further analyzed by a custom Python (v. 3.12.8) script in Jupyter Notebook using the modules *os* and *re* and the packages *matplotlib, numpy, pandas* and *scipy*.

Using the Python script, effects of antidromic stimulation on MN5 spike times during in-flight spiking behavior were either analyzed independent or dependent of stimulus occurrence within current phase of MN5 spiking (*cf*. Fig. 1E vs. Fig. 2A). Due to minute intrinsic fluctuations in MN5 interspike intervals during tonic spiking, stimulations that were administered in regular intervals (3, 5 or 10 s) occurred stochastically at different phases of MN5 spiking.

For each stimulus, its relative occurrence within the current MN5 spike cycle, as well as the effect of the stimulation on the subsequent spike timing were normalized to the *current (expected) average interspike interval (ISI) duration*: *i*.*e*., the mean over the last five MN5 ISI durations prior to stimulation.

Stimulation of a single additional MF spike *resets* the current MN5 spike cycle by starting a new spike cycle of approximately *current average ISI duration, i*.*e*. the ISI from stimulated spike to next orthodromic spike (*reset ISI, Fig. 1D*). Our analysis mainly focused on the effects of single pulse and pulse-train stimulations on the relative duration of the *reset ISI* and *post-reset ISI*: *i*.*e*., the duration of the *reset ISI* or *post-reset ISI* relative to the *current average ISI* duration.

To calculate the compensatory effect that the reset ISI and post-reset ISI durations pose to counteract instantaneous increases in firing rate that stem from single evoked spikes during tonic firing (Fig. 1F), we first calculated for each single pulse stimulation within each recording of Canton S. the time point at which the last unperturbed ISI prior to stimulation reaches phase 0.5. From this timepoint we counted the spikes within the one second leading up to this timepoint (*cf*. Fig. 1F: “pre-stim”) and the subsequent second starting at this timepoint (*cf*. Fig. 1F: “+ stim comp”). The first second thus representing unperturbed firing and the second second representing the subsequent second including the evoked spike and possible compensatory changes in spike frequency. We also calculated an expected spike count over one second that would result from single evoked spikes if no compensatory effect would occur, by taking the count of the second of unperturbed firing and adding one (*cf*. Fig. 1F: “+ stim exp”). All counts of unperturbed, expected and compensated firing for all stimulations occurring within each of the 23 recorded animals were averaged together respectively as median and are plotted in Fig. 1F.

### Whole cell patch clamp electrophysiology

Before dissection, adult 2-day old male w1118 (control) or HCN (Ih) excision mutant *Drosophila* were cooled down on a cooling plate and the red fluorescent lipophilic dye DiI (Thermo Fisher, D3899) was placed inside DLM fibers 5 or 6 with sharp minuten pins. DiI diffuses inside the plasma membrane, ultimately labeling the somata of those DLM MNs that are innervating the muscle fiber, DiI was placed in. After this procedure, flies were left in fly vials with food at room temperature for 2 hours before dissection. Then legs and wings were removed, and the animals were pinned dorsal side up in a Sylgard 184 (Farnell, Cat# 101697) coated 35 mm plastic dish with 0.1 mm minuten pins (Ehlert & Partner, Entomologie-Bedarf, Cat# 4141110). Dissection was done in normal *Drosophila* saline of the following composition [mM]: NaCl 128, KCl 2, CaCl_2_ 1.8, MgCl_2_ 4, HEPES 5, Sucrose 35, pH was adjusted to 7.24 with 1 N NaOH, osmolality was adjusted to 300 mOsM kg^-1^ with sucrose if necessary. DLM motoneurons (MNs) were exposed by cutting along the dorsal midline all the way to the head. Wing muscles were spread laterally and held down with minuten pins. Guts and esophagus were removed, the head was cut. After rinsing thoroughly with normal saline, the specimen was then mounted on an upright fixed stage Zeiss Axio Examiner A1 fluorescence microscope and viewed with a Zeiss W Plan Apochromat 40x/1.0 DIC VIS-R M27 water dipping lens. The MN5 soma is located at the dorsal surface of the mesothoracic neuromere of the ventral nerve cord. The somatic membrane was made accessible for patch clamp by removing the ganglionic sheath and removing debris using 1% Protease XIV from *Streptomyces griseus* (Sigma-Aldrich, P5147) in normal saline applied through a patch pipette with a manually broken tip. Whole cell patch clamp recordings in current clamp mode were conducted in normal saline (composition above). Patch pipettes were pulled from borosilicate glass capillaries without a filament (World Precision Instruments, PG52151-4) with a P-10 vertical electrode puller (Narishige), filled with internal patch solution of the following composition [mM]: 140 mM K-gluconate, 2 mM Mg-ATP, 2 mM MgCl_2_, 11 mM EGTA, 10 mM HEPES. pH was adjusted to 7.24 with 1 N KOH, osmolality was adjusted to 300 mOsM kg^−1^ with glucose if necessary. Patch pipette tip resistance was between 5 and 6 MΩ with these solutions. Recordings were conducted with an Axopatch 200B patch clamp amplifier (Molecular Devices) in current clamp mode, data were filtered at 5 kHz through a low pass Bessel filter, digitized at 20 kHz with an analog/digital converter (Digidata 1440, Molecular Devices). Output gain was 10x. After giga seal formation (> 5 GΩ), whole cell configuration was achieved by application of brief negative pressure to the patch pipette, rupturing the cell membrane. Only recordings with a holding current below -100 pA to hold a membrane potential of -70 mV as tested in voltage clamp mode were accepted for analysis. Series resistance as read from the respective dial on the amplifier did not exceed 10 MΩ and was corrected by ∼40%. Then mode was switched to current clamp, and the resting membrane potential was determined. To elicit tonic firing for frequency/input current curves (F/I curves), square current injections of 400 ms duration were applied with amplitudes between 0 and 0.9 nA in 0.1 nA increments. The resulting firing frequency was determined and plotted against input current amplitude.

### Immunohistochemistry of HCN channels

For cell type specific label of HCN channels in DLM MNs, we used the FLP-Tag method (Fendl et al. 2020). Briefly, an inverted GFP-containing cassette with likewise inverted splice acceptor and splice donor flanked by FRT sites can be flipped in the presence of flipase and may re-insert with a certain likelihood in the correct reading orientation. If paired with cell type specific expression of flipase, this results in GFP-tagged HCN channels only in the targeted cells. We expressed UAS-FLP (Bloomington # 4539, RRID:BDSC_4539) under the control of a DLM MN specific Split GAL4 driver w[*]; P{y[+t7.7] w[+mC]=R23H06-p65.AD}attP40; P{y[+t7.7] w[+mC]=R30A07-GAL4.DBD}attP2 (Bloomington #602182, Hürkey et al. 2023) in an Ih FLP Tag (HCN FLP-Tag) background. y[1] w[*]; Mi{FlpTag.2}Ih[MI12136-FT.2-X] (Bloomington # 92146, RRID:BDSC_92146, Fendl et al. 2020). For immunocytochemistry, flies were briefly anesthetized on ice, legs and wings were cut, and the fly was pinned dorsal side up in a sylgard coated plastic dish as for patch clamp electrophysiology. After dissection, the specimen was fixed with 4% paraformaldehyde in 0.1 M PBS (PBS) for 45 minutes, rinsed 5x with PBS and then washed 4×20 minutes with PBS at room temperature. This was followed by 4×20 minutes washes with 0.5% PBS-TritonX-100 (PBS-Tx). Then α-GFP FluotagX4-Abberior Star RED nanobody (Nanotag Biotechnologies, Göttingen, Germany, Cat# N0304-AbRED-L) was applied at a concentration of 1:500 in 0.2% PBS-Tx for 2 hours at room temperature in the dark, rocking. Preparations were then rinsed 5x with PBS and then washed 4×20 minutes with PBS followed by mounting in methylsalicylate. Specimen were covered with high precision cover slips (175 µm ± 5 µm) and sealed with clear nail polish.

HCN^GFP^ in DLM MNs was visualized with a Leica TSC SP8 confocal laser scanning microscope with a helium neon laser with 4% laser power at 633 nm excitation wavelength. Fluorescence was detected between 640 nm and 700 nm with a gallium arsenide hybrid detector with a gain of 40 %. Scanning speed was 600 Hz, voxel size was 80×80×300 nm, image depth was 8 bit (TIFF). Images were done with LasX software and CorelDraw 2025.

### Statistical Analysis

Statistical analysis was performed in GraphPad PRISM v.10.4.2. Data groups were tested for gaussian distribution by Shapiro-Wilk tests. For comparisons between groups containing non-normally distributed data or groups with sample sizes below n=10 non-parametric tests were used. Test for statistical significance are described in the respective results section. Statistical significance was defined as *p < 0.05, **p<0.01, ***p<0.001, ****p<0.0001.

